# Accelign: a GPU-based Library for Accelerating Pairwise Sequence Alignment

**DOI:** 10.64898/2025.12.17.694868

**Authors:** Felix Kallenborn, Fawaz Dabbaghie, Martin Steinegger, Bertil Schmidt

**Author notes:** Contributing authors.

## Abstract

**Background:** The continually increasing volume of sequence data results in a growing demand for fast implementations of core algorithms. Computation of pairwise alignments based on dynamic programming is an important part in many bioinformatics pipelines and a major contributor to overall runtime due to the associated quadratic time complexity. This motivates the need for a library of efficient implementations on modern GPUs for a variety of alignment algorithms for different types of sequence data including DNA, RNA, and proteins.

**Results:** Accelign is a library of accelerated pairwise sequence alignment algorithms for CUDA-enabled GPUs. Its parallelization strategy is based on a common wavefront design that can be adapted to support a variety of dynamic programming algorithms: local, global, and semi-global alignment of genomic and protein sequences with a variety of commonly used scoring schemes supporting one-to-one, one-to-many or all-to-all pairwise sequence alignments. This leads to a peak performance between 16.1 TCUPS and 9.1 TCUPS for computing optimal global alignment scores with linear gaps and affine gap penalties on a single RTX PRO 6000 Blackwell GPU, respectively. In addition, our library demonstrates significant speedups in several real-world case studies over prior CPU-based (SeqAn, Parasail, BSalign) and GPU-based libraries (ADEPT, GASAL2), and can even outperform highly customized algorithms (WFA-GPU, CUDASW++4.0). Furthermore, the performance of our approach scales linearly with the number of employed GPUs, which makes it feasible to exploit multi-GPU nodes for increased processing speeds.

**Conclusion:** Accelign provides significant speedups for commonly used pairwise alignment algorithms compared to prior implementations. It is freely available at https://github.com/fkallen/Accelign.

## Background

During the last decade we have witnessed a tremendous increase in the volume of data generated in the life sciences, especially propelled by the rapid progress of highthroughput sequencing technologies. Corresponding datasets typically contain large amounts of short reads (of a few hundred base-pairs in length for Illumina platforms), long reads (varying between a few thousands to hundreds of thousand base-pairs in length for Nanopore-based sequencers), or protein sequence collections (varying between a few and tens of thousands of amino acids).

Corresponding pipelines frequently use dynamic programming (DP) algorithms to compute batches of pairwise alignments with typical examples including local, global, or semi-global sequence alignments for DNA/RNA read mapping or protein homology search. However, the associated time complexity proportional to the product of alignment target lengths often makes them major contributors to overall runtimes. Modern bioinformatics tools therefore often require their highly efficient implementation. This motivates the need for solutions that can accelerate a variety of these algorithms on modern parallel computing platforms which can keep pace with the increasing volume of sequence data.

Consequently, a number of parallelized implementations have been developed for computing pairwise sequence alignments on a variety of architectures including CPUs [1–7], GPUs [8–20], FPGAs [21–25], PIM (Processing in Memory) [26], and even customized ASICs [27–29]. They typically target two types of application scenarios:

1. computing a single pairwise alignment of two very long DNA sequences (typically whole genomes or chromosomes), and
2. computing batches of different pairwise alignments of DNA/RNA or protein sequences of shorter length.

Widely adopted parallelization schemes are *inter-sequence* (computes a number of independent alignment tasks in parallel), *intra-sequence* (computes the DP matrix of a single alignment in parallel), and their *hybrid* combination.

There are several CPU-based libraries that provide multi-threaded and SIMD vectorized implementations of a variety of alignment algorithms including Seqan [1], SSW [5], Parasail [4], and BSalign [7]. In comparison, existing GPU-based alignment libraries are often limited in their functionality; e.g. ADEPT [14] only support local alignment of both DNA and protein sequences while GASAL2 [13] is limited to DNA sequences. Although, NVBIO [15] provides more functionality it has not been updated since 2015 and thus does not support modern GPU architectures. In addition, there are a number of more recent specialized GPU-based implementations. For example, WFA-GPU [30] is an algorithm for global alignment of DNA sequences under a certain type of scoring scheme.

We have recently presented CUDASW++4.0 [16] and MMSeqs2-GPU [17] for protein sequence database search based on computing local pairwise alignments. While CUDASW++4.0 is based on the Smith-Waterman for computing optimal alignments with affine gap penalties, MMSeqs2-GPU additionally uses a gapless pre-filter heuristic. We have shown that our novel parallelization strategy based on warp intrinsics for fast inter-thread communication and minimization of overall executed instructions can outperform prior approaches and achieve close-to-peak performance on modern GPU architectures. While this method is highly efficient, it has so far been limited to the use-case of protein homology search.

In this paper, we extend our parallelization strategy for accelerating a variety of commonly used pairwise alignment algorithms in genomics and proteomics. They are integrated in our new Accelign (<u>Acce</u>lerate and <u>align</u>) library, which contains the following features:

- **Sequence data types:** Accelign supports various alphabets such as those for DNA, RNA, and protein.
- **Alignment classes:** Accelerated kernels for local, global, and semi-global alignment are supported. Note that our kernels perform score-only computation but no traceback. However, for local alignment and semi-global alignment the start and the end-position can be output in addition to the score.
- **Scoring schemes:** We provide optimized kernels for linear and affine gap penalties. In addition arbitrary nulceotide or amino acid substitution matrices can be used or alternatively position-specific scoring matrices (PSSMs).
- **Application scenarios:** We support one-to-one alignments between two given sets of sequences, one-to-many alignments between a query and a set of sequences, and all-to-all alignments between two sets of sequences.
- **Multi-GPU:** Higher speeds can be achieved by using multiple GPUs connected to the same host by distributing the utilized datasets.
- **APIs:** The library is written CUDA C++ and we provide interfaces for its usage with C++.

We initially evaluate the theoretical peak performance of our various kernels using simulated datasets on different GPU architectures in terms of CUPS (<u>C</u>ell <u>U</u>pdates <u>P</u>er <u>S</u>econd). Subsequently, we evaluate the runtime performance of Accelign using real-world datasets consisting of Illumina reads and protein sequence databases. Our performance comparison shows that Accelign consistently outperforms other GPU-based libraries (GASAL2, ADEPT) and CPU-based libraries (SeqAn, Parasail, BSalign). In addition, it is faster than state-of-the-art specialized GPU-based implementations such as WFA-GPU for global alignment of Illumina reads and CUDASW++4.0 for protein database scanning. Even higher speeds can be achieved by using multiple GPUs connected to the same host with close-to-linear scaling. Last, we show the potential of an exact GPU-accelerated aligner in the context of read mapping for evolutionary divergent sequences.

Table 1 compares the features of all GPU-based pairwise alignment implementations considered in our performance evaluation. To summarize, Accelign not only performs best in our tests but also provides the highest flexibility.

**Table 1.**
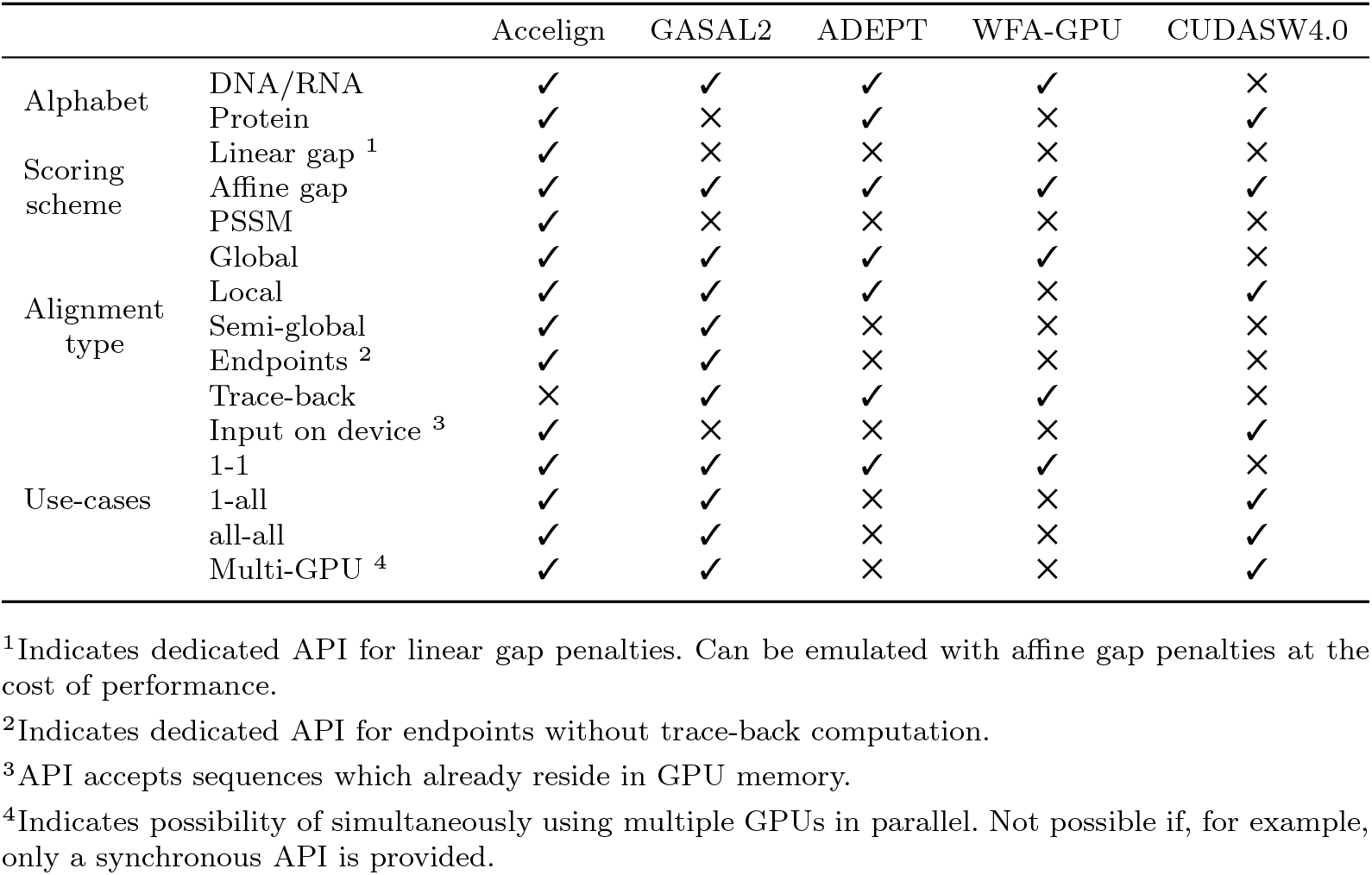
Feature comparison of all tested GPU-based pairwise alignment implementations.

### Pairwise Alignment Computation

Pairwise sequence alignment considers two sequences *S* = (*s*_1_*s*_2_ … *s*_*n*_) of length *n* and *Q* = (*q*_1_*q*_2_ … *q*_*m*_) of length *m* over the alphabet Σ. For each pair (*s*_*i*_, *q*_*j*_) of characters we decide if these characters should be aligned or if a gap should be inserted and record a corresponding score in a DP matrix *H* using the recurrence relation:

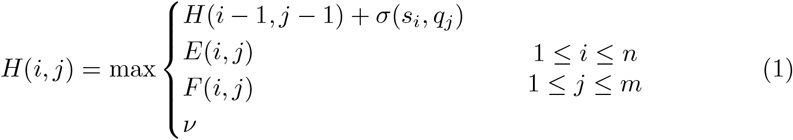

where *σ* is a substitution function over Σ × Σ that determines the score of aligning two characters. The parameter *ν* is needed to distinguish between local and global alignments as we will explain below. *H*(*i, j*) stores the optimal alignment score of the prefixes (*s*_1_ … *s*_*i*_) and (*q*_1_ … *q*_*j*_) with respect to the utilized scoring scheme.

In case of a *linear gap penalty α* we set:

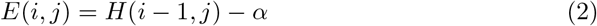

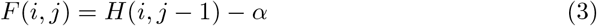

*Affine gaps* penalize a gap of length *k* by *α* + (*k* − 1) · *β* where *α* is the cost of the first gap (opening) and *β* the cost of each subsequent gap (extension):

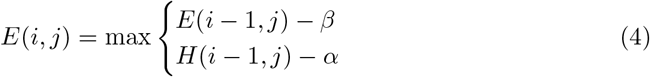

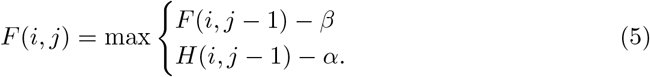

Initialization of the first rows and columns of *H, E*, and *F* – as well as in what cell(s) to look for the optimal score – depends on whether the alignment shall be *global, local*, or *semi-global*.

*Local alignments* compute an alignment with the highest score starting at any position (*i, j*) and ending at any other position (*k, l*) in *H* with *i* ≤ *k* ≤ *n, j* ≤ *l* ≤ *m*. The parameter *ν* in Eq. 1 is set to 0. The DP matrices are initialized by *H*(0, 0) = *H*(*i*, 0) = *H*(0, *j*) = 0, *E*(0, 0) = *F* (0, 0) = *F* (*i*, 0) = *E*(0, *j*) = −∞, *E*(*i*, 0) = −*α* − (*i* − 1) · *β*, and *F* (0, *j*) = −*α* − (*j* − 1) · *β*.

*Global alignments* always start at position (0, 0) and end at position (*n, m*). The optimal global alignment score is also stored in cell *H*(*m, n*). The variable *ν* in Eq. 1 is set to −∞. The DP matrices are initialized as: *H*(0, 0) = 0, *H*(*i*, 0) = *E*(*i*, 0) = −*α* − *β*(*i* − 1), *H*(0, *j*) = *F* (0, *j*) = −*α* − *β*(*j* − 1), *E*(0, 0) = *E*(0, *j*) = *F* (0, 0) = *F* (*i*, 0) = −∞.

*Semi-global alignments* do not penalize gaps at the beginning and the end of the alignment. This leads to the same specifications as for local alignment with two exceptions: (i) *ν* = −∞, and (ii) the optimal score is determined as the maximal value in the last row or column of *H*.

Figure 1 illustrates the conceptual layout and dependency relationship between cells of *H*. Note that score-only computations of all these variants can be performed in linear space O(min{*n, m*}) and quadratic time O(*n* · *m*).

**Fig. 1.**
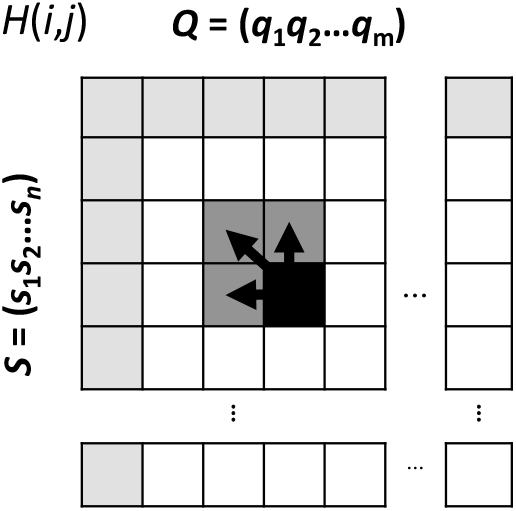
Conceptual layout and dependency relationships of the DP matrix *H*. Light gray cells are initialization cells and dark gray indicates the ancestral subproblems of the currently active cell (black).

### Position Specific Scoring Matrices

A PSSM (Position Specific Scoring Matrix) – also known as a PWM (position weight matrix) or a profile – is a frequently used representation in which substitution scores are given separately for each position in a protein or DNA profile; i.e., it features one row per symbol of the considered alphabet Σ and one column per position of the profile. PSSMs are often obtained from multiple sequences alignments (MSAs) and are for example often used for homology search in protein sequence databases or for DNA motif discovery.

Typical tasks require computing pairwise alignments between a sequence *S* = (*s*_1_*s*_2_ … *s*_*n*_) of length *n* and a PSSM *Q* represented by a matrix of size |Σ| × *m*; i.e., the value *Q*[*s*_*i*_, *j*] represents the likelihood of observing *s*_*i*_ ∈ Σ at position 1 ≤ *j* ≤ *m*. The optimal alignment score can be computed by replacing the lookup *σ*(*s*_*i*_, *q*_*j*_) in Eq. 1 by the lookup *Q*[*s*_*i*_, *j*].

Note that storing the PSSM generally requires more data than the substitution function. Thus, fast memory access to the PSSM is key to achieve high efficiency for computing pairwise alignments between sequences and PSSMs.

### Use Case Scenarios

Bioinformatics applications that require large amounts of pairwise alignments typically consider two given sets ℛ = {*R*_1_, …, *R*_*N*_} and 𝒯 = {*T*_1_, …, *T*_*M*_} containing *N* and *M* sequences, respectively.

### One-to-One

For this use case holds *N* = *M* and refers to computing an alignment between *R*_*i*_ and *T*_*i*_ for all 1 ≤ *i* ≤ *N* resulting in a total of *N* pairwise alignment computations. This is a common task in various applications including read mapping. We will consider such alignments of Illumina reads in our first case study.

### One-to-All

This corresponds to the special case where one of the sets contains only a single sequence; i.e., *N* = 1. Thus, *R*_1_ is aligned to all sequences in 𝒯 resulting in a total of *M* pairwise alignments. This task occurs in database search applications such as protein homology searches, which we consider in our second and third case study.

### All-to-All

This use case refers to the task of aligning each sequence in ℛ to each sequence in 𝒯 resulting in a total of *N* × *M* alignments. It can be expressed as multiple one-to-all alignments.

### Features of CUDA-enabled GPUs

CUDA kernels are executed using a number of independent thread blocks. Each thread block is mapped onto exactly one streaming multiprocessor (SM) and consists of a number of warps. All 32 threads within a warp are executed in lock-step fashion (SIMT).

CUDA-enabled GPUs contain several types of memory: large but high latency global memory and fast but small on-chip shared and constant memory. Nevertheless, the fastest way to access data is through usage of the thread-local register file.

Modern GPUs provide instructions for warp-level collectives in order to efficiently support communication of data stored in registers between threads within a warp without the need for accessing global or shared memory.

A crucial feature of our approach is the usage of warp shuffles for low latency communication and minimization of memory traffic. In particular, we take advantage of the warp-level collectives __`shfl _down_sync()` and `__shfl_up_sync()` e.g. the intra-warp communication operation `R1 = __shfl_up _sync(0xFFFFFFFF, R0, 1, 32);` moves the contents of register `R0` in thread *i* within each warp to register `R1` in thread *i* + 1 for 0 ≤ *i <* 31.

Recent GPU generations have incrementally included key hardware features that can be used to improve the performance of DP algorithms.

- `Half2:` Support of half-precision floating point arithmetic instructions that allow for the simultaneous execution of one 32-bit operation on two half-precision values. This allows for computing two values of the DP recurrence relation within a single register, effectively doubling the throughput. Ampere and subsequent generations enable the necessary `half2 min/max` comparison function present in pairwise alignment recurrence relations.
- `Max3:` Hopper introduced a new half2 set of instructions to perform the min/max operation of three input values (`vhmnmx`).
- `DPX:` Inclusion of DPX instructions to support fast computation of typical DP recurrence relations with integer-based arithmetic has been introduced with the Hopper architecture. This includes a set of functions that enable finding a max value of three inputs (`vimax3`), as well as fused addition with max operations (`viaddmax`). The instructions support any combination of min/max, sub/add for 16-bit or 32-bit integer parameters, with optional ReLU (clamping to zero).

In the following section we will describe the necessary algorithmic changes to achieve high performance using these features for designing our new GPU-based library.

### Implementation

#### Core Parallel Algorithm

We base our parallelization scheme on computing an independent alignment per (sub)warp – a group of synchronized threads executed in lockstep that can efficiently communicate by means of warp shuffles with typical sizes of 4, 8, 16, or 32 threads — called a **cooperative group (CG)** in the following. Multiple CGs within a thread block handle different alignments independently, while the threads within a CG compute DP matrix cell values of a single alignment in cooperative fashion. Note that according to Eqs. 1, 4, 5 each DP cell depends on its left, upper, and upper-left neighbor (see Fig. 1), which means that (parallel) computation of DP cells has to follow this topological order. This dependency relationship holds for all alignment algorithms considered in our library and allows for the computation of cells along a minor diagonal of the DP matrix in parallel (also called a **wavefront**).

The core algorithm of Accelign for computing a single pairwise alignment within a CG in parallel is based on a common wavefront design and works as follows. Consider a CG consisting *p* threads *T*_0_, …, *T*_*p*−1_ with *p* ∈ {4, 8, 16, 32} and the two sequences *S* = (*s*_1_*s*_2_ … *s*_*n*_) of length *n* and *Q* = (*q*_1_*q*_2_ … *q*_*m*_) of length *m* = *p* · *k*. We assign *k* columns of the DP matrix to be calculated to each thread. Computation proceeds along a **wavefront** in *n* + *p* iterations. In iteration *i*, thread *T*_*t*_ computes *k* adjacent cells of the DP matrix row *i* − *t*. All DP cells of the current and previous iteration are stored in thread-local registers. The required access of thread *T*_*t*_ to the rightmost value of thread *T*_*t*−1_ computed in the previous iteration is accomplished by using the lowlatency warp shuffle instruction `__shfl_up _sync()`. The described mapping strategy is illustrated in Fig. 2.

**Fig. 2.**
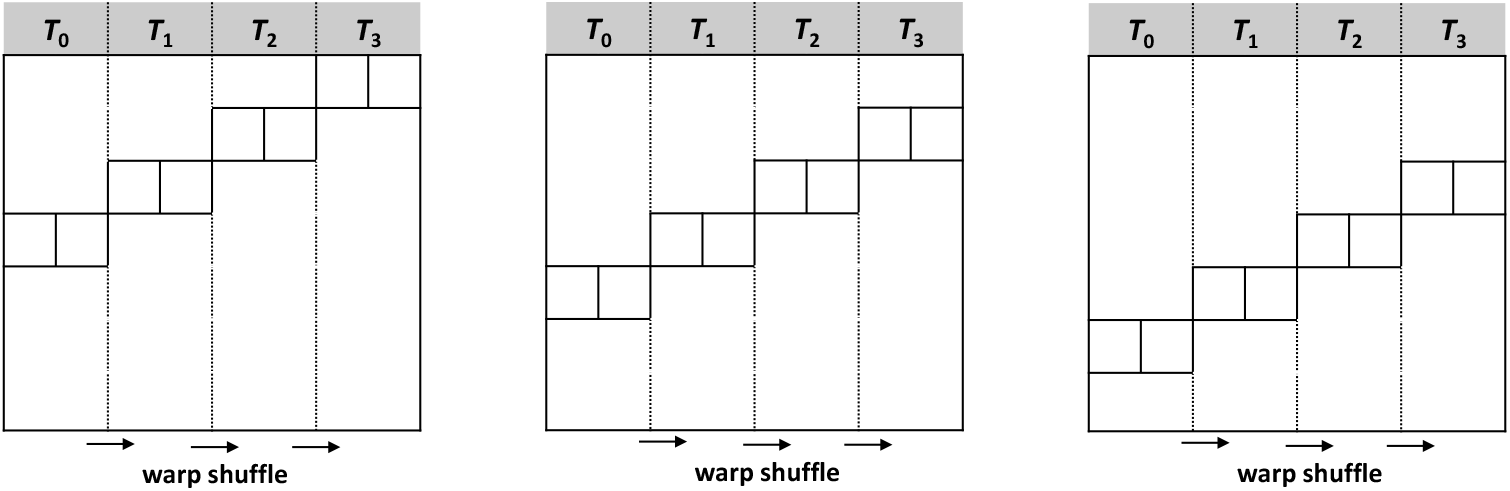
Illustration of three iterations of our wavefront DP matrix computation with *p* = 4 threads and *k* = 2 columns per thread. Each thread computes *k* = 2 new matrix cells in each iteration using the values of the antecedent row and an additional value from the second antecedent row which are all stored in thread-local registers. Threads communicate values in the rightmost of their *k* columns by a warp shuffle operations (arrows).

Listing 1 shows pseudocode of a GPU device function for the described core algorithm which is called by each CG to compute a different global alignment with a linear gap penalty using `float` 32-bit arithmetic. Template parameters are used to define the number of DP matrix columns assigned to each thread (`k`) and the CG size (`p`). Thus, by changing the definitions of *p* and *k*, kernels tailored for different sequence sizes can be generated. Prior to the wavefront loop, we transfer *Q* to thread-local registers (`QReg[]`) and initialize the DP matrix values stored in each thread in the registers `M[], M_left, M_diag` in `initDP()`. `processDP()` computes *k* DP cells per thread in each wavefront iteration (highlighted cells in Figure 2) using only threadlocal in-register computation and shared memory lookups to the PSSM or substitution function (as will be explained in the corresponding subsection below). The rightmost current DP cell of each thread stored `M[k]` is then communicated to Register `M_left` of the neighboring thread using a warp shuffle-up instruction. Every *p* iteration steps, we load one value from *S* per thread to Register `SReg cache`, while Register `SReg` always stores the character from *S* needed in each iteration step. A warp shuffle-up and a warp shuffle-down update `SReg_cache` and `SReg`, respectively, with characters from neighboring threads.

In the following we will describe how this basic kernel can be extended to implement the various pairwise alignment variants considered in Accelign.

**Listing 1.**
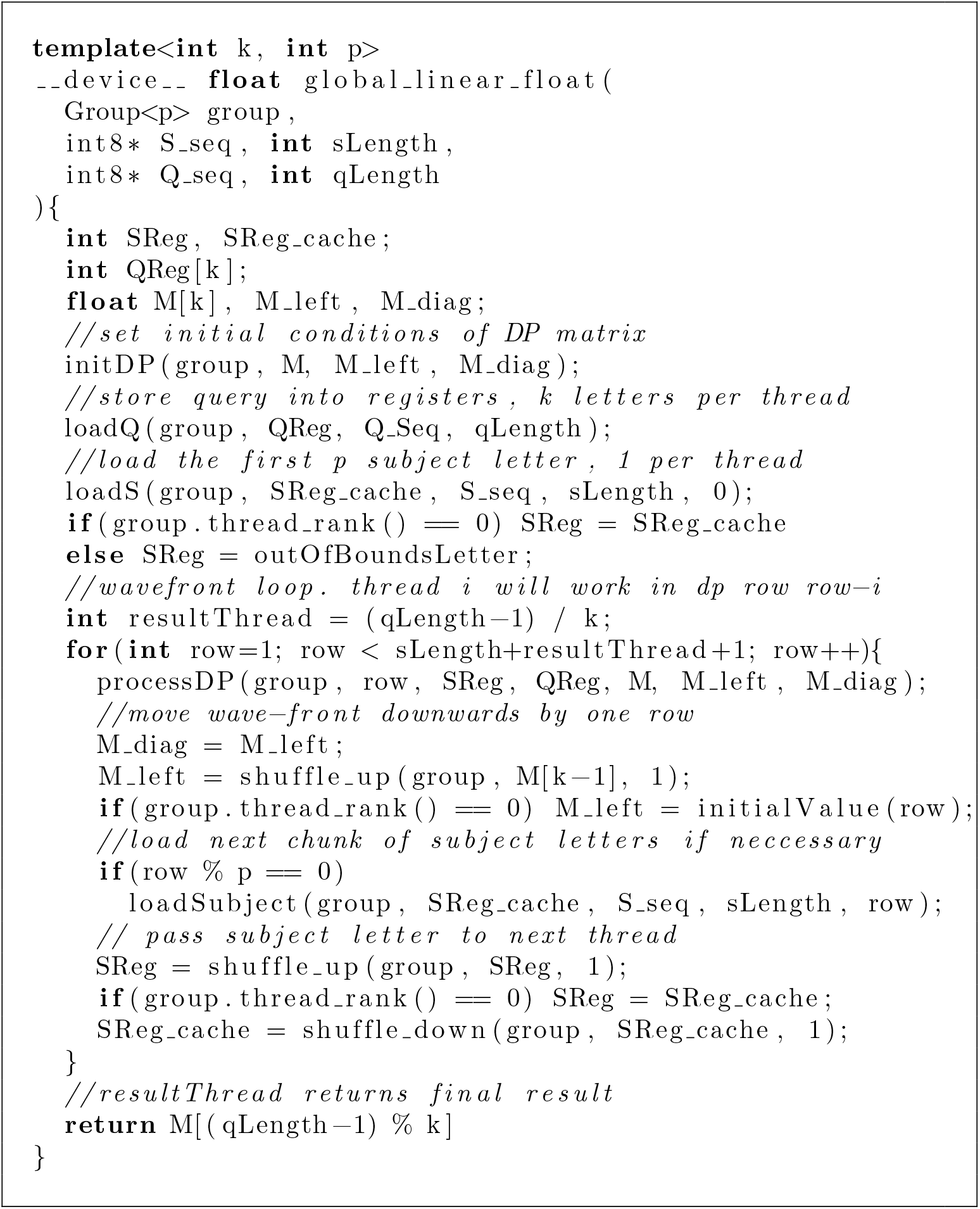
Pseudo-code of our core parallel wavefront algorithm for linear global alignment using float computation.

### Kernel Variants

All computational kernels provided by our library follow the general code structure presented in Listing 1. We use optimized functions tailored towards the computation of global alignment, local alignment, and semi-global alignment, with different penalty schemes and datatypes. In the following, we will describe implementation details for each algorithm. We begin with observations which hold true for all kernels.

For each algorithm type, we provide a so called *AlignmentState* which represents the wavefront. It encapsulates the DP matrix registers and contains the logic to initialize and compute the DP matrix cells. This allows for a simple switch between optimized implementations for linear gap penalty and affine gap penalty, as the gap penalty scheme is independent from the movement of the wavefront within the DP matrix.

The efficiency of Accelign relies on storing and computing DP cells in thread-local registers. However, thread-local arrays are typically only materialized in registers if the compiler can compute all accessing indices at compile-time. We achieve this by manual unrolling or automatic loop unrolling using `#pragma unroll`.

The DP cells can be computed using different datatypes. Our library supports scalar types like 32-bit `float` and `int`, as well as vector types with two components like packed 16-bit types `half2` and `short2`. Our mathematical operations are specialized for each supported datatype. For example, the operation `max(a+b,c)` which is common to alignment computations is implemented as written for floats, but uses a specialized function `_viaddmax s32(a,b,c)` for integers, which maps to a single hardware DPX instruction on supporting GPU architectures like NVIDIA Hopper or Blackwell GPUs. We do not provide specialized kernels for DNA or Proteins per se. Instead, our alignment kernels can simply be specialized for a specific alphabet size |Σ| which is passed as a template argument. All input characters must be encoded in the range [0, | Σ| − 1]. Our library does not pose additional restrictions to the encoding. Internally, the alphabet size |Σ^*′*^| = |Σ| + 1 is used. It contains an additional element representing an out-of-bounds character, which is returned by out-of-bounds accesses to input sequences. Out-of-bounds accesses occur when the wavefront is not fully contained in the DP matrix. When `half2` or `short2` is used, each CG computes two independent alignments in the upper and lower 16 bits, respectively. Internally, the corresponding two query sequences and the two subject sequences will be zipped together producing a new sequence, respectively, encoded in the range [0, |Σ^*′* 2^| − 1]. Next, we present details specific to each supported alignment algorithm.

### Global Alignment

The optimal score of a global pairwise alignment is always stored in the bottom-right cell of the DP matrix. In our wavefront, this cell can in theory be covered by any thread, but more importantly it appears at position `M[(qLength-1)%k]`, which is an array index not known at compile-time. To ensure that the compiler can still place the array in registers, we explicitly copy the array to local memory after computation is complete to access it via dynamic indexing without restrictions. A different challenge regarding the output arises when using the packed 16-bit types. In general, input sequences can be of different length. The global alignment score of any sequence must be output as soon as its cell has been computed. Otherwise, the value may be overwritten in the next row. Therefore, we need to split the alignment loop into two parts when dealing with `half2` or `short2`: First, compute all rows required for the shorter of the two sequences, and output the respective score. Then, continue until the longer sequence is fully processed.

### Local Alignment

An optimal local alignment score computation returns the maximum observed value. During wavefront computation, each thread thus keeps track of its observed maximum score. After the DP matrix is fully processed, a group-wide parallel reduction is performed to find the total maximum across the group. It is then returned. Note that for global alignments the computation of out-of-bounds DP cells cannot affect the final score as the thread which holds the result is fixed for a given alignment calculation. However, this may not be the case for a local alignment. The out-of-bounds computation could change the per-thread observed maximum, and the total maximum could be located in any thread. To avoid this issue, we use a substitution score ≤0 for any out-of-bounds positions. Thus, the per-thread observed maximum can never change by out-of-bounds DP cell calculations. This approach is also beneficial when using `half2` or `short2`. Since computation can arbitrarily extend beyond the DP matrix without changing the final result, the alignment loop does not need to be split into two parts. A single loop with maximum number of DP rows is sufficient.

Compared to a global alignment, a local alignment performs an additional clamping to 0. Although CUDA provides specialized (DPX) functions for `int` and `short2` to compute max(max(*a* + *b, c*), 0), it is only hardware-accelerated since the Hopper architecture, and no such function exists for `float` and `half2`. However, the operation max(*a* + *b*, 0) can be implemented for `half2` using a fused multiply-add with integrated relu, `__hfma2 relu(a,half2(1,1),b)`, which maps to a single hardware instruction on all GPU architectures since Ampere. The local alignment recurrence can be reformulated using max(*a* + *b*, 0) which allows the usage of the specialized function for `half2` which saves one instruction in the inner loop. For example, with linear gap penalty, the instruction sequence

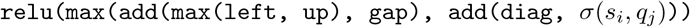

can be rewritten as

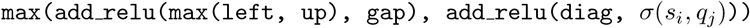

This is based on the observation that if the score produced by one direction is negative, it can never become the new maximum as the maximum is at least 0. Thus, negative values can be clamped to 0. Then, since all the directions now produce a non-negative score, the final relu operation can be omitted. We also apply a similar code transformation for affine gap penalty.

### Semi-global Alignment

Computing a semi-global alignment requires tracking the maximum in the last column of the DP matrix. As explained previously, the thread ID and the per-thread column ID for the last column of the DP matrix depends on the query length and is not know at compile-time. To avoid conditional branches in the inner loop, we unconditionally track the maximum of all columns at the cost of higher register pressure. The wavefront loop needs to be split into multiple parts. Part A includes all iterations where the wavefront covers the initialization row of the DP matrix. Part C includes all iterations covering the last row of the DP matrix. In between, there is part B which contains all other iterations. Note that for short sequences part B can be empty, and parts A and C may overlap. Part A is necessary to explicitly initialize the first row of the DP matrix to 0 for threads other than the first thread. In contrast, local alignment does not require explicit initialization because 0 is implicitly generated via the relu operations. Part C enables maximum tracking within the last row. Similar to local alignment, this produces a per-thread local maximum which is transformed into a total maximum via group-wide parallel reduction. After the computations are complete, the thread which covers the last DP matrix column computes the maximum between the total row maximum and the maximum observed in the last DP matrix column to produce the final result.

For global alignment with packed 16-bit types, we split the wavefront loop into two part to account for the potentially different matrix sizes of the two sequences. This is further complicated for semi-global alignment, leading to five parts. Part A for the initialization, part C and part E for tracking the last row in the two DP matrices, and parts B and D for the remaining iterations.

### PSSM and Substitution Matrix Lookups

Common to all of our alignment variations is the need to compare a position in the subject with a position in the query, producing a substitution score *σ*(*s*_*i*_, *q*_*j*_). Such substitution scores are commonly stored in memory as substitution matrix or PSSM. Note that simple scoring approaches could also compare the two letters explicitly to produce a *matchscore* or *mismatchscore*.

Our alignments support both a substitution matrix and a PSSM. The simple case of explicit comparison could be modeled by a substitution matrix which stores *matchscore* on its main diagonal, and *mismatchscore* otherwise (see Fig. 3(a)). Note that for a given query and substitution matrix it is trivial to compute a corresponding PSSM.

**Fig. 3.**
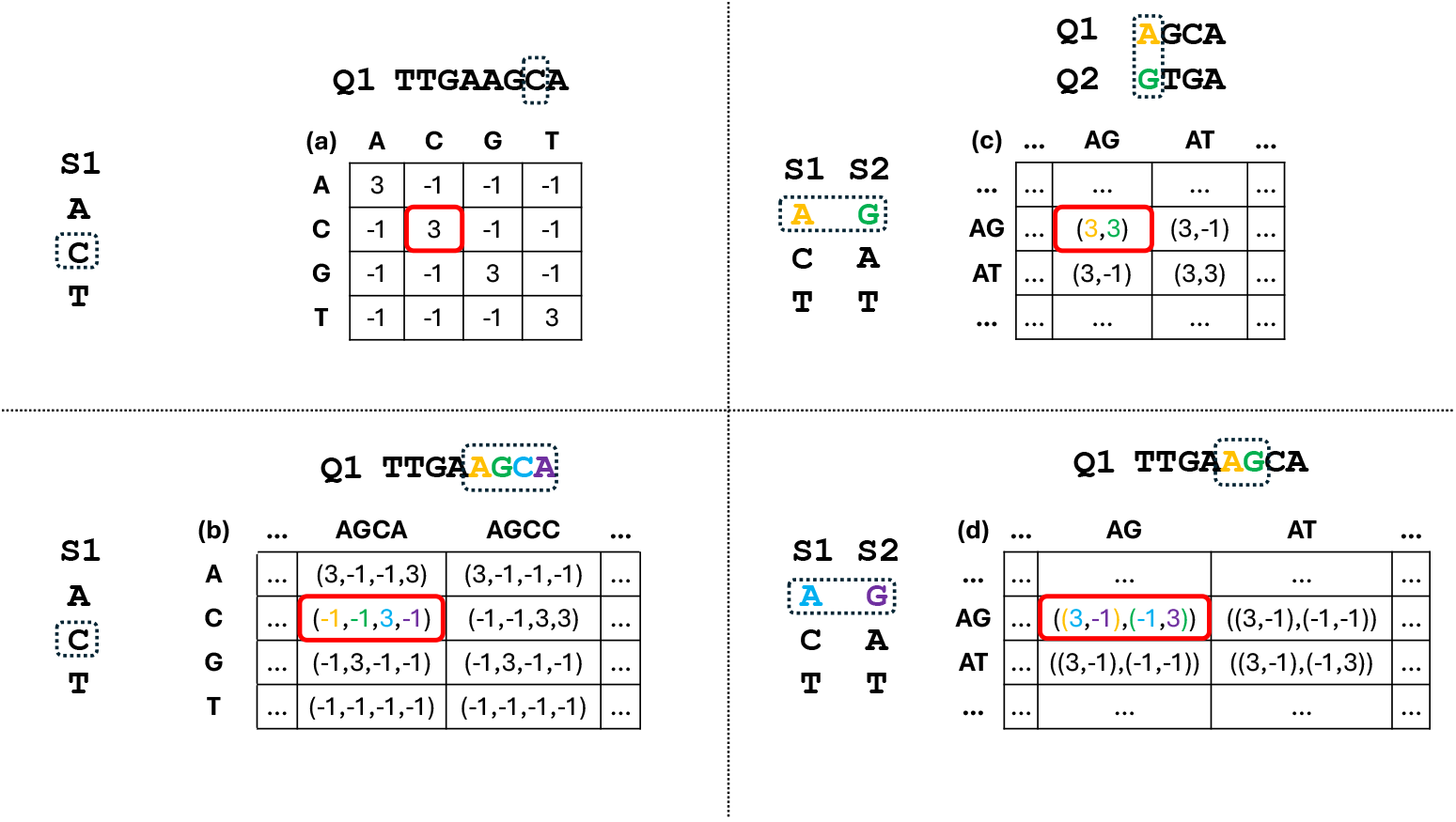
Substitution scores of substrings in dashed boxes are given by red boxes within a substitution matrix. Numbers and letters are colored to indicate corresponding scores. The specified input substitution matrix will be converted into one of many internal layouts. These include: (a) Traditional layout. (b) Substitution matrix for an alignment using scalar (e.g. float) computations. All combinations of four letters are stored in one dimension to load substitution scores of four consecutive letters in one sequence. (c) Layout for vector (e.g. half2) computations. Two zipped sequences (S1,S2) are aligned against two zipped sequences (Q1,Q2). Substitution matrix stores the element-wise comparison result. (d) Layout for vector (e.g. half2) computations. Two zipped sequences (S1,S2) are aligned against one sequence (Q1). A pair of zipped letters is compared against two consecutive letters of the single sequence.

In our implementation we use a level of abstraction which allows for simple integration into the main algorithm. The substitution scores are returned by a so called *SubstitutionScoreProvider*. It provides functions which take *s*_*i*_ as input and return the substitution score for one or multiple columns covered by a thread. We implemented different substitution score providers for PSSM and substitution matrix, and for different score datatypes.

When calling an alignment kernel, the scoring matrix or PSSM is initially stored in (high latency) global memory. As a first step before computing any alignments, the substitution scores are copied into faster shared memory which can be accessed by all threads within a thread block. Current generations of CUDA-enabled GPUs typically provide at least 99KB of shared memory per thread block. To efficiently access shared memory, one needs to use an appropriate access pattern. Shared memory is accessed by a full warp; i.e., 32 consecutive threads. It is organized into 32 4-byte memory banks which can operate in parallel. Each shared memory address is served by a specific fixed memory bank. The *i*-th four-byte value in the shared memory address space corresponds to bank *b* = *i* mod 32. A warp-wide access to shared memory is decomposed by hardware into 128-byte-wide transactions. A shared memory bank conflict occurs if multiple different addresses which map to the same memory bank are part of the same transaction. As the memory bank cannot serve multiple different addresses at the same time, the transaction needs to be replayed as often as required to serve all addresses of the transaction. If possible, one should minimize the number of shared memory bank conflicts as they reduce the memory throughput.

After covering the fundamentals, we will now explain the implementations of the different score providers in more detail.

### Substitution Matrix

Accelign allows the use of a substitution matrix for all alignment application scenarios. It is passed as an integer matrix of size |Σ| × |Σ|. Internally, we can choose between multiple matrix shapes with respect to

- matrix dimensions,
- layout in shared memory, and
- data type.

The specific choice depends on the use case and is a trade-off between

- shared memory access speed,
- shared memory size, and
- executed instruction count.

First, consider the case where a scalar score type is used for DP cell computations and the matrix shape is |Σ^*′*^| × |Σ^*′*^| storing the same scalar type.

Since all input sequences are properly encoded as integers, we can use those integers as index into the substitution matrix to return the corresponding substitution value. Note that this requires to keep the query letters in registers to avoid reloading them in each iteration of the wavefront loop. Considering a whole CG, the access pattern for the substitution matrix is random access, as each thread within the group effectively operates with different subject/query value pairs. In particular, this implies random access to shared memory within a warp. Thus, we are susceptible to bank conflicts as the memory bank requested by each thread is effectively random. However, in the case of DNA alignments with Σ = {*A, C, G, T, N*} this is not a problem. The substitution matrix has a memory size of just 144 bytes (when using `float` or `int`), where the last 6 · 4 = 24 bytes are only ever accessed for out-of-bounds subject letters. This leaves 30 elements to be accessed frequently, each assigned to a different bank. Thus, for the majority of accesses there will be no bank conflicts. For larger alphabets such as those used for proteins, however, this no longer holds true.

Note that one shared memory lookup per DP cell is performed in the case described above. This could be reduced by using wider memory loads, loading multiple substitution scores per instruction. Yet, due to the random access pattern it is unclear if the additionally loaded elements are necessary for subsequent DP calculations. However, by increasing the size of the substitution matrix in shared memory it is possible to utilize wider memory loads without loading unused data. This can be achieved by using a row and/or column for each possible combination of two or four elements of the alphabet (see Fig. 3(b)). The substitution scores of multiple consecutive sequence positions can then be obtained by a single look-up using the respective combination of letters as the index into the inflated substitution matrix. This concept is particularly important for DP calculations using vector types like `half2` which require two substitution scores for the different alignments in each vector lane. Examples of different matrix layouts with their use-case are shown in Figure 3.

In addition to the different matrix shapes, a second degree of freedom is the choice of a substitution score datatype. To reduce the amount of shared memory used by the substitution matrix, one could consider using a narrower element datatype such as 16-bit floats instead of 32-bit floats. This can prove beneficial for performance even if it introduces an additional instruction to convert from the narrower type to the type used for DP cell calculation. For example, we observed a case where the performance was limited by an oversubscribed memory input/output (MIO) queue. The optimal matrix layout for this case stored all combinations of 4 letters and used 16-bit floats. While this did not reduce the number of instructions (4 32-bit loads vs 1 64-bit load + 4 conversion instructions), it shifted work to a lesser utilized hardware pipeline which in turn led to increased performance.

### PSSM

A PSSM typically requires significantly more memory than a substitution matrix because its size depends not only on the alphabet size, but also on the query length, or rather *p* · *k*, the maximum number of columns which fit the CG. Due to the potentially high shared memory requirements when using a PSSM, we only support PSSM in the case of one-to-many alignments which requires to store only a single PSSM in shared memory. From a performance perspective a PSSM allows for the best possible shared memory access pattern. The reason is two-fold:

1. Each thread in a group accesses a fixed distinct set of *k* consecutive columns of the PSSM; i.e., thread *i* will only access columns [*i* · *k*, (*i*+1) · *k*). Therefore, it is possible to reorder the PSSM columns in memory to enable bank-conflict-free accesses.
2. Since a thread requires multiple consecutive elements within the same PSSM row, 16-byte-wide memory load instructions can be used to reduce instruction count in the innermost loop. Those fetch four required 32-bit substitution scores in a single instruction.

Additionally, in contrast to substitution matrix, since the mapping of per-thread DP matrix columns to shared memory PSSM columns is fixed, the actual query letters are not longer required which reduces register pressure.

One particular challenge is to provide bank-conflict-free accesses for kernel configurations where the total amount of bytes loaded per CG in one instruction is less than the shared memory transaction size of 128 bytes. This case arises for example with a group size of 4 and 16-byte loads. Then, the loads of the next CG, which will access the same PSSM columns, but from potentially different rows, are included in the same 128-byte transaction. This can lead to a bank conflict. Our solution is to replicate the PSSM in shared memory as often as needed to ensure that groups which are part of the same transaction access distinct memory banks, leading to conflict-free access. In the described example with four threads and 16-byte loads, we utilize two PSSMs which are interleaved such that the *g*-th group only accesses memory banks 0 − 15 when *g* is even, and only accesses banks 16 − 31 when *g* is odd. Figure 4 provides an example of a conflict-free PSSM layout in shared memory.

**Fig. 4.**
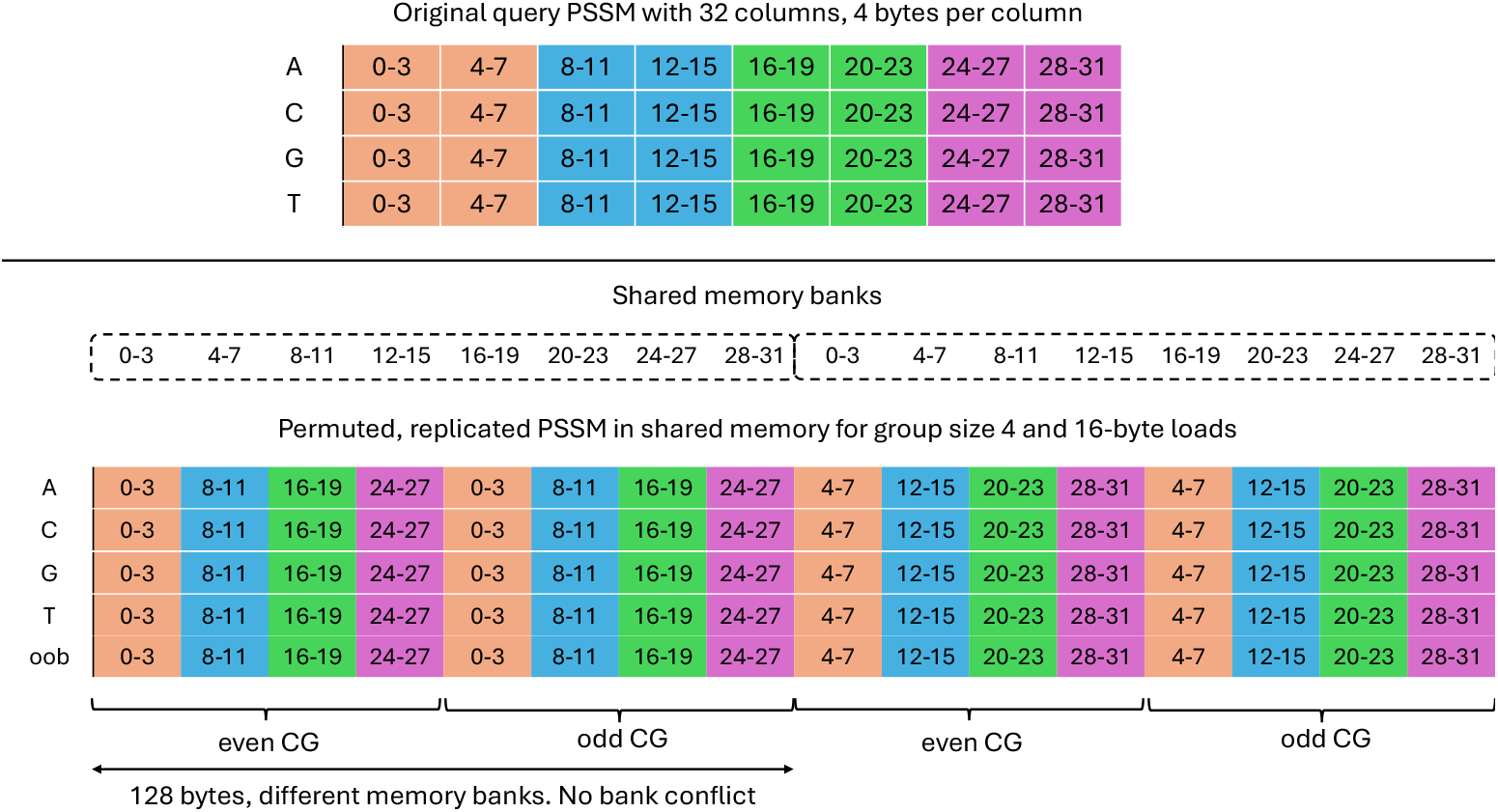
A PSSM with 32 columns should be aligned by CGs with 4 threads, and 8 columns per thread, i.e. *p* = 4, *k* = 8. Different colors indicate different threads within a CG. Since threads in a CG execute concurrently, PSSM columns which are accessed at the same time should be stored consecutively which requires a permutation of columns. By interleaving two copies of a permuted PSSM, two CGs can be served by a single conflict-free 128-byte transaction which uses distinct memory banks.

### Memory usage and its limitations

Conceptually, our library supports any alphabet size. However, in practice the alphabet size is limited by the available shared memory to hold the PSSM or substitution matrix.

Especially for the `half2` or `short2` types, where memory grows at least quadratically with alphabet size a limit is quickly reached. To enable the usage of those types with alphabet sizes typical for bioinformatics applications, e.g. protein alphabet, we have also implemented low-memory score providers. Here, the value type stored in the data structure are only `half` or `short` instead of `half2` and `short2`. Additionally, we no longer use the zipped alphabet to access the data. Instead we perform a separate lookup for each of the two sequences and pack the two values into a `half2` or `short2` for calculation. Using this low-memory approach the shared memory size only grows linearly with alphabet size for `half2` and at `short2` the cost of additional instructions.

With 99 kilobyte of shared memory per block and |Σ| = 25, the low memory variant for `half2` and `short2` is necessary for both the PSSM and the one-to-one substitution matrix for most matrix tile sizes. The maximum alphabet size for which the lowmemory `half2` PSSM variant is able to process a query sequence of length 1024 with 99 kilobytes shared memory is 48 when using 2-byte storage types. A low-memory `half2` substitution matrix can be used with alphabet sizes up to at least 128.

Note that the simplest approach to bypass shared memory size limitations would be to keep all data in global memory. While functional, this would be prohibitively slow. Our library therefore does not offer this option.

### Finding start and end positions of alignments

At present time, Accelign is designed to compute alignment scores, but cannot compute the full traceback paths. However, for local alignment and semi-global alignment we provide additional kernels that are able to identify the end coordinates of the alignment path within the DP matrix, and optionally computing the start coordinates as well. Functionality of computing start/end positions is limited to alignments with substitution matrix and datatypes `float` and `int`. PSSM and packed 16-bit types are currently not supported.

To solve the task of finding the end coordinate the algorithms proceed as usual, with the addition of keeping track of the coordinate of the current observed maximum score in the DP matrix for local alignment (or in the last row and last column in the case of semi-global alignment). While updating the observed maximum can be done in a single instruction, explicit conditionals are required to determine if the observed maximum changed, and if yes, to update the corresponding cell coordinates. This adds additional instructions in the innermost loop which reduces the performance compared to score-only alignments.

Our approach to find the start coordinates is the computation of an optimal alignment of the reversed input sequences. Let *S* and *Q* be the alignment input sequences with |*S*| = *n* and |*Q*| = *m*. We first use the aforementioned approach to find the optimal alignment score *O* and its DP cell coordinates (*e*_*s*_, *e*_*q*_) which corresponds to the end position of the alignment. This gives us the information that the optimal alignment is produced by substrings *S*[1: *e*_*s*_] and *Q*[1: *e*_*q*_]. Next, we compute an alignment with end position of the reversed substrings *S*^*′*^ = *rev*(*S*[1: *e*_*s*_]) and *Q*^*′*^ = *rev*(*Q*[1: *e*_*q*_]). This gives DP cell coordinates 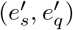 of the end of the reverse alignment, which can be converted into start coordinates (*b*_*s*_, *b*_*q*_) of the original forward alignment. Using these start and end coordinates, external libraries could be used to compute a global alignment between *S*[*b*_*s*_: *e*_*s*_] and *Q*[*b*_*q*_: *e*_*q*_] to obtain a traceback.

As an optimization to our approach, we use the already computed alignment score from the forward alignment to short-cut the reverse substring alignment. As soon as the optimal alignment score is observed in the DP matrix, we can conclude that the end has been found as there cannot be a better score, and terminate.

### Partitioning for long sequences

Our kernels can be configured to support different query lengths via parameters *p* and *k*. However, *p* and *k* cannot be increased indefinitely due to hardware limitations. First, warp shuffles can only be used for *p* ≤ 32. Second, the *k* DP matrix cells need to be kept in per-thread registers, whose number is limited. Excessive register spilling to slow global memory will hinder performance. Our solution to support queries of length *l > p* · *k* is matrix tiling. We define the *tile size* as *z* = *p* · *k*. Then the DP matrix can be partitioned into ⌈*l/z*⌉ many matrix tiles with up to *z* columns per tile. A single CG then processes the matrix tiles in order. When the CG processes any matrix tile which is not the last tile, it writes the computed DP cells of the last column of the tile to temporary storage in global memory. This enables processing of the successor tile. The saved DP cells are required to serve the diagonal and left dependency for the first column the successor tile, and are loaded on demand.

The size of the required temporary storage per CG depends on the maximum subject length max(*l*_*s*_) to be processed, the group size *p*, and the number of bytes to be written per DP cell. Specifically, our current implementations requires (max(*l*_*s*_) + *p*) * *cellsize* bytes per CG. For alignments with linear gap penalty, 4-byte elements are used. The temporary storage for alignments with affine gap penalty requires 8 bytes per cell. Note that we require separate temporary storage for each CG. To limit the total size of the temporary storage, a grid-strided for-loop is used to enable processing of multiple alignments per CG in a round-robin manner. This allows us to limit the number of CGs to the maximum number of CGs that can run concurrently on the GPU.

### Accuracy of data types for computing alignment scores

By using SIMD-like instructions to operate on two 16-bit quantities packed in a 32-bit value, it is theoretically possible to achieve up to two-fold speedup. However, using 16-bit numbers for sequence alignment can have draw-backs in terms of accuracy. Regardless of the used data type, only integer scores are represented. The 16-bit half datatype can represent all exact integer values *x* where −2048 ≤ *x* ≤ 2048. For 16-bit shorts the range is −32768 ≤ *x* ≤ 32767. Integer values outside of these ranges are rounded towards zero for half. For example, the value 2049 is rounded to 2048. In the case of shorts, no values outside the given range are representable, and signed integer overflow/underflow may occur during calculations, which is undefined behavior in the `C++` standard.

Accelign does not check if a value cannot be represented during computations. Such checks would cancel out the performance advantage of 16-bit types due to increased instruction count. We recommend using the traditional 32-bit types instead if a wider range of values needs to be supported. Nevertheless, for local alignments a simple post-processing step can identify a score as exact. Let *x* be the final 16-bit alignment score, and *b* the largest entry in the substitution matrix. For all cases where *x <* 2048 (`half2`) or *x* + *b* ≤ 32767 (`short2`), *x* is the exact result.

## Results

We evaluated the performance of Accelign on the following machines:

**S1** (Blackwell) AMD EPYC 7713P CPU (64 physical cores, 128 logical cores), 512 GB RAM, 2 Nvidia RTX PRO 6000 Blackwell GPU (2 × 96 GB VRAM).

**S2** (Ada) Dual-socket Intel Xeon Gold 6548Y+ (2 × 32 physical cores, 120 logical cores), 1 TB RAM, 2 Nvidia L40S GPU (32 GB VRAM).

**S3** (Ampere) AMD EPYC 7713P CPU (64 physical cores, 128 logical cores), 512 GB RAM, Nvidia A100 GPU (80 GB VRAM).

**S4** (Grace Hopper) ARM Neoverse-V2 CPU (72 physical cores, 72 logical cores), 480 GB RAM, Nvidia GH200 (96 GB VRAM).

The primary system used for our evaluation are S1 and S4. The other systems are used for comparison between different GPU generations on a synthetic benchmark. CUDA 13.0 and gcc 9.3.0 are used as GPU and host compiler, respectively.

As a preliminary step, on each system we conducted a grid search over a subset of possible alignment configuration parameters for alphabet sizes 5 (DNA/RNA) and 25 (Protein). The results were used to derive a set of GPU-specific matrix tile sizes, thread configurations, and PSSM/substitution matrix configurations for best performance.

Next, based on the obtained configurations we developed a high-level convenience API on top of our low-level kernel-based API. Given a batch of sequence pairs to align, the high-level API is able to select a suitable alignment configuration based on the maximum query length observed within the batch. For queries with a length ≤512 a single matrix tile is used whose size is the best fit to the query length. For longer queries, we switch to multi-tile processing with a tile size of up to 1024. The selected tile size requires the least number of tiles and performs the fewest out-of-bounds computations. Substitution scores are restricted to the range [− 15, 15] since some configurations make use of the 8-bit floating point type `__nv_fp8_e4m3`. This still allows us to use the common BLOSUM substitution matrices for protein alignments. It is possible to prevent usage of this type, which increases the supported range to [−2048, 2048].

In the following, we demonstrate the performance of our library in different synthetic and real-world case studies. For one-to-all alignments the PSSM approach is used. In the case of one-to-one alignments a substitution matrix is used instead. Unless stated otherwise, the high-level API is used.

### Performance scaling with sequence length

In this synthetic benchmark we processed a batch of simulated sequences of equal lengths. Sequence length was varied between 32 and 65536 in steps of powers-of-two. The batchsize was limited to 4 gigabytes worth of data, and 2^26^ sequences. We gathered performance data for all of our supported algorithms, alphabet sizes 5 and 25, and different data types used for computation.

We measure performance in terms of matrix cell updates that are performed per second; for example, TCUPS (Trillions of Cell Updates Per Second) as

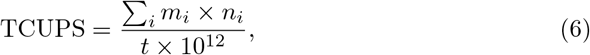

where *t* is the runtime in seconds and *m*_*i*_ and *n*_*i*_ are the lengths of the aligned sequences. Here, we only give a summary of our findings in Figure 5. The full set of benchmark results can be found in the Supplementary File.

**Fig. 5.**
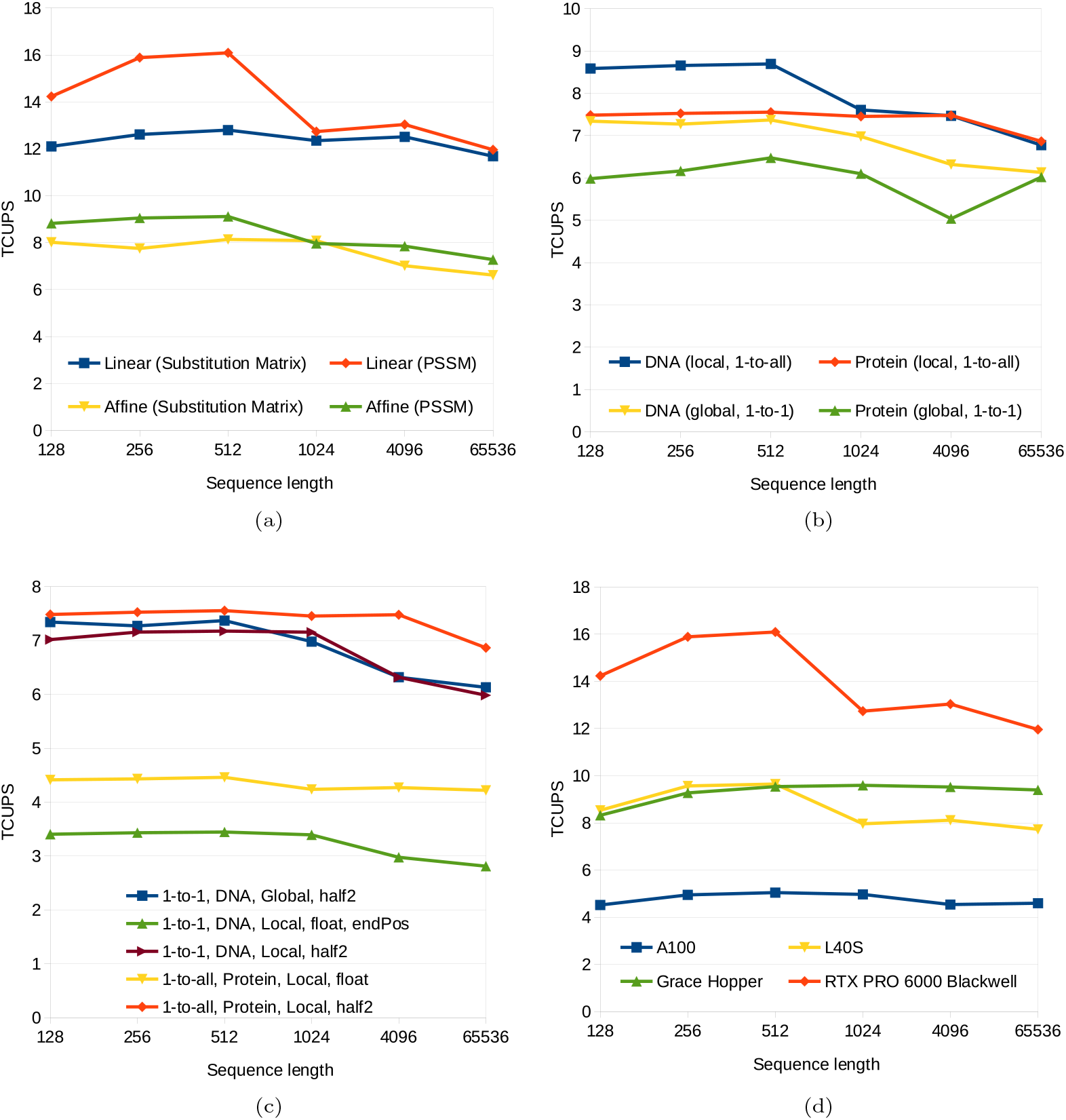
(a) Performance comparison between substitution matrix and PSSM using a single GPU on S1. A global alignment of DNA was used with 16-bit half2 computations. (b) Performance comparison between alphabets for DNA/RNA and proteins using 16-bit half2 computations with affine gap penalty using a single GPU on S1. (c) Synthetic benchmark of different kernels with affine gap penalty used in our real-world case studies on a single GPU of S1. (d) Performance comparison of different GPU generations computing a 1-to-all DNA global alignment with linear gap penalty, half2 computations, and PSSM.

First, we motivate the choice of PSSM for one-to-all alignments by taking a look at the performance differences between PSSM and substitution matrix. Substitution matrix performance was measured using the kernel-level API. Figure 5a shows the achieved performance on single GPU of S1 across the different sequence lengths for both approaches for a global alignment of DNA sequences using `half2`. Calculating a global alignment with linear gap penalty requires the fewest computations per DP cell. Any inefficiencies in shared memory accesses and instruction count will thus be noticeable. Indeed, for short queries with a length up to 512 the PSSM variant is up to 29% faster. For long queries, performance is impacted by global memory accesses due to matrix tiling, hiding the benefits of PSSM. Still, a speedup of 5% can be observed. On the other hand, alignments with an affine gap penalty come with a greater compute-to-memory-access-ratio. Here, using a PSSM improves the performance by up to 16% (11%) on short (long) queries.

Next, Figure 5b shows the performance differences between the two alphabet sizes when using `half2` and affine gap penalties. In this case, alignment kernels are more efficient when applied to DNA sequences. This is because of the aforementioned challenges in shared memory usage. Additional instructions are required for the protein alphabet with low-memory variants of PSSM and substitution matrix. When 32-bit floats are used instead (not present in the figure), one-to-all alignments show similar performance on both alphabets because the same PSSM layouts and storage types can be used. On the other hand, one-to-one alignments are faster on the smaller alphabet because a substitution matrix layout which supports a wider memory access size can be employed.

In our real-world case studies, a number of different alignment algorithms were used. Figure 5c highlights the corresponding performance. The half2 version of a one-to-all local affine alignment on proteins is around 70% faster than the float version. For one-to-one alignments, computing the alignment end position in addition to the optimal score costs up to 25% performance. One peculiar observation is that for an affine semi-global alignment the multi-tile kernel is noticeably faster than the single-tile variant. We did not investigate this further.

Last, Figure 5d provides a performance comparison between the latest Blackwell generation and prior CUDA-enabled GPU architectures. We have chosen a DNA one-to-all linear global alignment with `half2` because it is the kernel which achieves the highest TCUPS across all tested GPUs. On this kernel, RTX PRO 6000 Blackwell is between 2.6x and 3.4x faster than A100, and between 1.5x and 1.9x faster than L40S, showing that performance increases with newer GPU generations.

### Case Studies

In the following case studies, we demonstrate the benefits of using our GPU-accelerated alignments with Accelign in three problem settings: DNA short-read alignment, Protein database searches, and HIV short-read mapping.

### Short-read alignment

We used Accelign to create a simple short read alignment tool. Given two fasta files with DNA sequences as input, it computes the global affine alignment score between the *i* − th sequence of each file. This task maps to the one-to-one case of our library with alphabet size 5. The program uses a three-stage pipeline with a dedicated CPU thread per stage. The first stage generates batches of sequences to be aligned. As part of a larger framework, this stage could involve other (pre-)processing steps. In our program, we simply parse the input files. The second stage receives such a batch from the first stage and aligns the contained sequences. This could be achieved by calling a CPU library or GPU library. In the case of GPU libraries, the typical workflow includes transferring the sequence data to the GPU, and subsequently executing a GPU kernel for alignment. Additional set up may be performed on the CPU. After the alignment kernel completes, the alignment scores are transferred back to the CPU. The last pipeline stage receives a batch of sequences and their corresponding alignment scores for further processing. Our benchmark tool creates an output file from the previously calculated alignment scores.

GPU alignment libraries often expect a list of CPU-side DNA strings as input and hide the required data transfers to the GPU from the user. While such an interface is simple to use, it has disadvantages in regards to performance. First, all sequence strings internally need to be copied into a contiguous staging buffer for efficient data transfer. Second, if the sequences were already present in GPU memory, they would first need to be copied to the CPU and transformed into strings before the alignment library could be called (copying them back to the GPU). Third, maintaining a large collection of separate strings may lead to memory fragmentation and poor data locality. In contrast, Accelign only provides the core alignment functions which expect the input data to already be present in GPU memory. Data transfer is explicitly managed by the user. Importantly, this allows us to accept the alignment input data in a contiguous data layout, which avoids creating an intermediate copy of the data. Thus, our first stage provides the sequence data in two formats; a list of separate strings for other APIs, and a contiguous buffer of sequence data for our API.

With above alignment framework set up, we measured the alignment performance of Accelign and compared it against other libraries. For this, we implemented the second pipeline stage for each library. Runtimes are only measured for this stage. We compared our library to the GPU-based implementations ADEPT, GASAL2, and WFA-GPU. In addition, we used Parasail 2.6.2, Seqan 3.4.0, and BSAlign 1.2.1 as CPU-based alignment libraries. The CPU benchmarks were performed on system S1 with 128 threads since these typically scale with the number of logical cores. System S4 was used for GPU benchmarks as it provides the fastest CPU-GPU interconnect, eliminating bottlenecks caused by data transfers.

A simple scoring scheme was used with gap opening = −3, gap extension and mismatch set to −1, and match score = 0. The match score was set to 0 because it is a precondition for WFA-GPU to work.

To generate the input data for our benchmark, we used BWA [31] to map Illumina human sequencing reads (accession number SRR1766553^1^) against reference HG002, and extracted the corresponding genome coordinates. The benchmark then attempts to align each read to its determined genomic subsection. In total, our benchmark dataset contains 10 million reads of length 148. The genomic subsections have minimum length of 103, a maximum length of 209, with an average of 148. The number of DP cells to be computed is approximately 219 · 10^9^.

Table 2 presents the runtimes for pipeline stage two. Since it is difficult to measure GPU-side activity accurately when transfers and GPU kernel launches are hidden behind the API, we used the NVIDIA Nsight Systems profiler with NVTX ranges to obtain durations of GPU activities as well as the total time spent on processing the sequence batches.

**Table 2.**
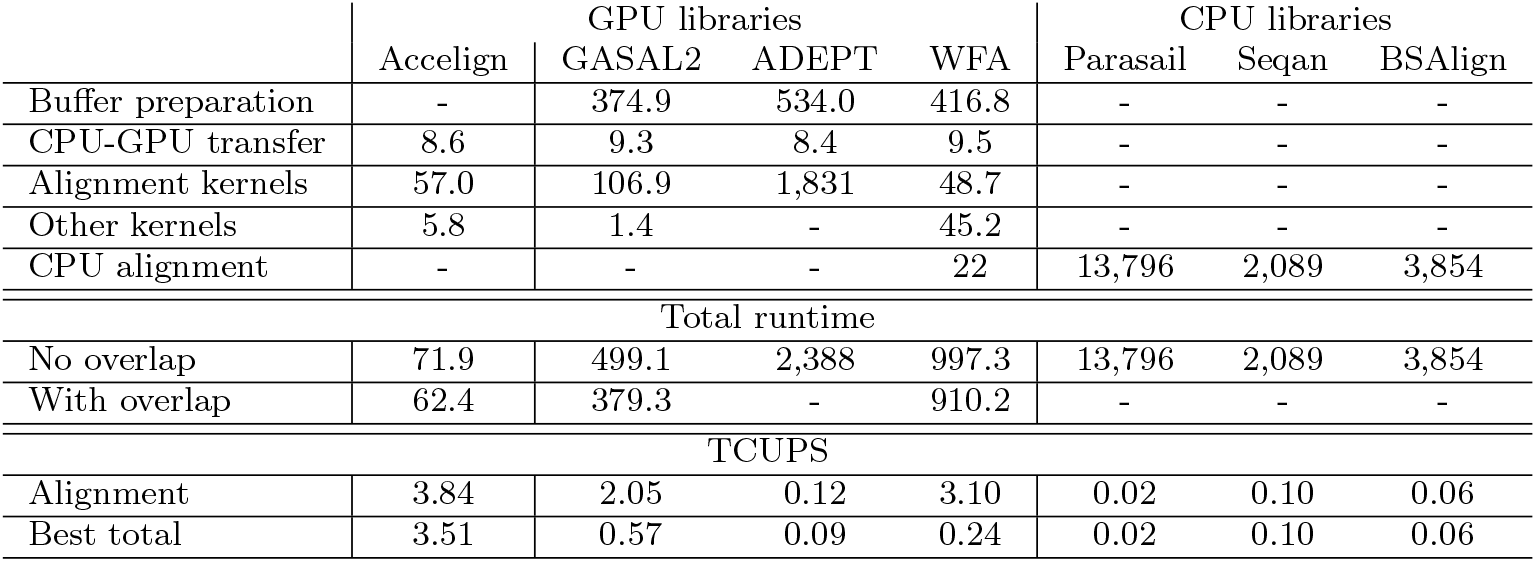
Runtimes (in milliseconds) and TCUPS for the second pipeline stage on system S4. Individual runtime contributions are only given for the (no overlap) case. Total runtime includes contributions not shown in the table. Accelign uses half2 computations.

Accelign achieves the fastest runtime for alignments which corresponds to 3.8 TCUPS. Compared to a synthetic peak performance benchmark, this is around 20% less than the maximum observed TCUPS. This can be explained by inefficiencies due to non-equal sequence lengths in the real-world data as opposed to the synthetic case. WFA-GPU has a fast alignment kernel as well but is unable to compute all alignment scores on the GPU due to algorithmic limitations. It needs to fall back to CPU-based alignments for those cases which reduces performance. ADEPT is outclassed by all other tested GPU libraries, and might be slower than fast CPU aligners on capable hardware.

The aggregation of sequence strings into a contiguous buffer takes at least 374 ms, which is the major contributor to the overall runtime of GASAL, ADEPT, and WFA-GPU. In addition, WFA-GPU is spending a significant amount of time in other steps (not shown in the table) due to additional CPU work which looks like initialization of other data. In contrast, Accelign can accept a contiguous sequence buffer as input, avoiding additional memory copies.

The next step involves transferring the data from CPU RAM to GPU VRAM. With observed fast transfer rates of up to 350 GB/s due to Grace Hopper’s NVLink interconnect, the data transfer does not pose a bottleneck compared to the duration of compute kernels. However, on systems with a traditional PCIe interconnect, for example PCIe 5.0 (∼ 50 GB/s), performance can be limited by data transfer rate, rather than compute performance.

The total pipeline performance using our library can further be improved by using CUDA streams to overlap PCIe transfers with GPU kernels, yielding a total of 3.51 TCUPS including CPU and GPU work as well as data transfers. CUDA streams are also supported by GASAL2 and WFA-GPU to overlap different stages within the pipeline.

### Protein Database Scanning

The second case study focuses on the topic of similarity search. Given a database of reference sequences and a query sequence, the goal is to find those reference sequences which are the most similar to the query sequence. One typical use-case is protein homology search as performed, for example, by BLASTP. Accelign supports database searches, for example via one-to-all alignment kernels utilizing a PSSM. A suitable alphabet size can be set to handle DNA/RNA, or proteins.

To benchmark our performance, we have set up the following experiment. The UniRef50 protein database (Release 2025 03 consisting of 20,080,454,799 amino acids in 70,198,728 sequences with largest (average) sequence length of 45,354 (282)) was used as reference. Similarity search was performed for a set of 20 real-world protein queries with sequence lengths varying between 144 and 5, 478. The employed alignment was a one-to-all PSSM local alignment with affine gap penalties (gap open = −11, gap extend = −1) and the BLOSUM62 substitution matrix (BLASTP default settings).

As a first step, we compute all alignments using fast packed 16-bit computations. This gives the exact alignment score for the majority of reference sequences. However, a small fraction of scores could be incorrect due to the limited resolution of 16-bit calculations. Thus, after 16-bit alignments we inspect the alignment scores and recompute the alignment score using 32-bit calculations if the 16-bit score is not flagged as exact.

Since Accelign is designed as a kernel-level API, its users may directly use the functions on multiple GPUs. We also demonstrate this capability in the context of the current benchmark. One approach to a multi-GPU database scan is to partition the database across the different GPUs, search the input query in the database partitions of all GPUs simultaneously, and merge the per-GPU result lists into a final result. In our case, we decided to partition the database such that each GPU computes a similar number of DP matrix cells. More advanced partitioning schemes may be used in real-world applications to achieve better load balancing. For example, it may be beneficial to have a similar number of alignments per GPU in addition to a similar number of matrix cells. In any case, our trivial partitioning is already sufficient to demonstrate the library capability and benefits of using multiple GPUs.

We compare the performance of Accelign to the GPU-based CUDASW++4.0 and the CPU-based BLASTP (version 2.17.0+) using 128 CPU threads on S1. For all programs, the reference database was first converted into a binary format suitable for processing. For CUDASW++4.0 and Accelign benchmarks, the resulting binary database fits completely into GPU memory. The duration of this pre-processing step and database transfer from CPU memory to GPU memory is excluded from our evaluation.

Table 3 reports the performance of CUDASW++4.0 and Accelign in terms of TCUPS for each of the 20 queries. For Accelign we are evaluating two configurations per query: high-level API (HL) and customized (C). The customized setting uses a configuration tailored towards the specific query length; e.g., the tilesize 464 = 16 · 29 was introduced which avoids out-of-bounds computations for query 4. Using a single GPU on S1 Accelign achieves an average performance 6.44 TCUPS with high-level API mode and 6.73 TCUPS in customized mode, outperforming CUDASW++4.0 (6.38 TCUPS on average) and BLASTP with 128 CPU threads (2.10 TCUPS on average). Furthermore, using both GPUs on S1, our library with a custom config achieves a speedup of 88%–96% over the single-GPU case, demonstrating efficient scaling to a multi-GPU setting.

**Table 3.** Database scan performance (TCUPS) with real protein queries. HL: indicates higher-level library implementation. C: indicates implementation with custom library config. BLASTP achieves an average of 2.10 TCUPS over all queries. Speedups are given over CUDASW4.

| Query | Length | CUDASW4 | Accelign |  |  | Speedup |  |
| --- | --- | --- | --- | --- | --- | --- | --- |
|  |  |  | HL | C | C (2 GPUs) | (HL) | (C) |
| 0 (P02232) | 144 | 5.62 | 6.27 | 6.27 | 11.81 | 1.12 | 1.11 |
| 1 (P05013) | 189 | 5.90 | 7.04 | 7.04 | 13.27 | 1.19 | 1.19 |
| 2 (P14942) | 222 | 6.00 | 7.17 | 7.16 | 13.96 | 1.20 | 1.19 |
| 3 (P07327) | 375 | 6.25 | 6.93 | 6.93 | 13.59 | 1.11 | 1.11 |
| 4 (P01008) | 464 | 6.32 | 6.59 | 7.07 | 13.85 | 1.04 | 1.12 |
| 5 (P03435) | 567 | 6.37 | 6.22 | 7.09 | 13.86 | 0.98 | 1.11 |
| 6 (P42357) | 657 | 6.41 | 5.92 | 6.73 | 12.64 | 0.92 | 1.05 |
| 7 (P21177) | 729 | 6.43 | 6.50 | 6.50 | 12.36 | 1.01 | 1.01 |
| 8 (Q38941) | 850 | 6.46 | 6.49 | 6.50 | 12.34 | 1.01 | 1.01 |
| 9 (P27895) | 1,000 | 6.48 | 6.85 | 6.85 | 12.99 | 1.06 | 1.06 |
| 10 (P07756) | 1,500 | 6.53 | 6.47 | 6.82 | 13.32 | 0.99 | 1.05 |
| 11 (P04775) | 2,005 | 6.55 | 6.61 | 6.83 | 13.35 | 1.01 | 1.04 |
| 12 (P19096) | 2,504 | 6.57 | 6.20 | 6.76 | 13.13 | 0.94 | 1.03 |
| 13 (P28167) | 3,005 | 6.58 | 6.58 | 6.81 | 13.31 | 1.00 | 1.03 |
| 14 (P0C6B8) | 3,564 | 6.59 | 6.55 | 6.84 | 13.24 | 0.99 | 1.04 |
| 15 (P20930) | 4,061 | 6.57 | 6.57 | 6.80 | 13.15 | 1.00 | 1.03 |
| 16 (P08519) | 4,548 | 6.56 | 5.86 | 6.66 | 12.72 | 0.89 | 1.01 |
| 17 (Q7TMA5) | 4,743 | 6.53 | 6.05 | 6.24 | 11.90 | 0.93 | 0.96 |
| 18 (P33450) | 5,147 | 6.47 | 6.17 | 6.14 | 11.70 | 0.95 | 0.95 |
| 19 (Q9UKN1) | 5,478 | 6.38 | 5.84 | 6.49 | 12.38 | 0.92 | 1.02 |

### HIV Read Mapping

Aligners such as BWA and Bowtie2 [32] do not construct a complete DP matrix between the two sequences aligned, but instead rely on finding short exact matches or seeds first, followed by local alignment extension from these seeds. This is computationally efficient as long as exact seeds can be found between the two sequences, which can be challenging for evolutionary divergent sequences. The Human Immunodeficiency Virus (HIV) is one of these challenging examples, with diversity reaching up to 20% within the same subtype and over 30% between different subtypes [33]. This higher diversity leads to fewer matched seeds and alignments.

To showcase that an optimal aligner can perform better in such biological cases, we simulated 500,000 short, single-end reads from an HIV Subtype O assembly (accession number KU168292) retrieved from the Los Alamos National Laboratory HIV database. We used the ART Illumina simulator [34] v.2.5.8 with the MiSeq Version 3 setting (`-ss MSv3 -l 250`) to produce the simulated sequences. We aligned the simulated sequences against the HIV reference genome HXB2 (accession number K03455) using both BWA-MEM v.0.7.19 and Accelign. For both aligners, we used a match score of 1, a mismatch penalty of 3, a gap opening penalty of 6, a gap extension penalty of 1 and a minimum score cutoff of 30. Accelign was able to align 179,301 sequences compared to BWA that aligned only 77,992 sequences; all sequences aligned by BWA were also aligned by Accelign. Figure 6(a) shows the coverage of both aligners over the HXB2 genome, with the depth calculated over 100-bp bins using mosdepth v.0.3.10. From the plot, we can see that Accelign was able to align many more sequences in the same regions and also align sequences to regions where BWA was not able to. Specifically, aligning sequences to the envelope (gp120) protein containing the highly variable V1-V5 loops, which are crucial in evading immunity [35]. Figure 6(b) plots the coverage counts for each gene in the reference; the counts were calculated using Bedtools version 2.31.1 with the `--counts` flag. This bar plot further highlights how Accelign was able to align many more sequences to genes, especially the envelope protein.

**Fig. 6.**
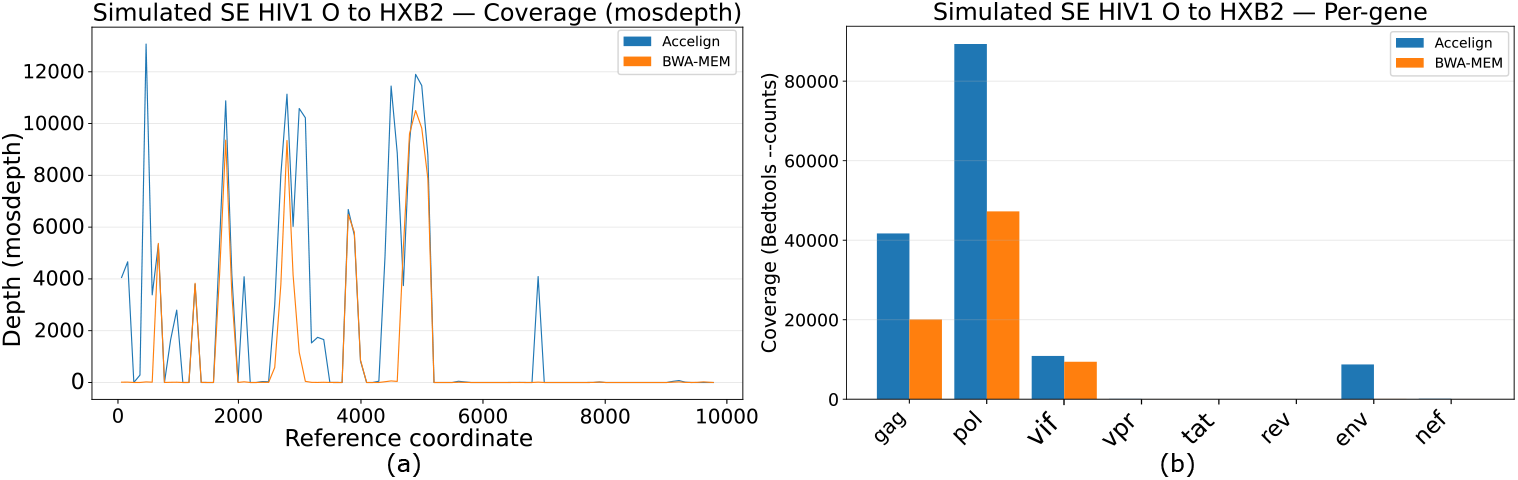
Results for the simulated single-end reads from the HIV1 O subtype aligned against the HXB2 reference using both Accelign and BWA. Panel (a) shows the average coverage over the reference genome for both Accelign (blue) and BWA (orange). Panel (b) shows the coverage counts for each gene in the HIV reference genome. Similar to (a), we see that Accelign was able to align many more sequences for certain genes, and align sequences against the envelope protein, where BWA was not able to.

We used the following workflow to compute the best mapping with Accelign. First, we employed a 16-bit score-only local affine alignment using a substitution matrix to align both the forward read and its reverse complement against the reference. With the given sequence lengths and scoring scheme, the optimal local alignment score is guaranteed to be less than 2048 which removes the need of re-computing scores in greater precision. Furthermore, orienting reads along the horizontal axis of the DP matrix allows to process the resulting tall matrix using a single matrix tile without temporary storage.

Next, for each read where at least one of the two directions reaches the minimum score cutoff, we identify the direction with the best score. While we do not provide traceback functionality in our library, it can nonetheless be used to speed up subsequent traceback computations. Specifically, for the identified set of best alignments, we recomputed the alignments with our kernel that finds the start and end positions of the traceback in addition to the alignment score. The resulting positions as well as alignment scores are subsequently transferred to the CPU. As a final step, Block Aligner [36] is used to compute the tracebacks of the sub-sequences given by the start and end positions.

On system S1 with 128 threads, BWA reports a processing time of 0.78 seconds (`M::mem_process_seqs`), whereas our implementation takes 0.5 seconds for GPU alignments and 0.2 seconds for CPU-sided tracebacks with Block Aligner, including all data transfers between CPU and GPU. To summarize, both programs require a similar amount of time for the core alignment step but BWA produces less mapped reads.

## Conclusion and Discussion

The continually increasing volume of sequence data in the life sciences results in a growing demand for high-throughput sequence processing pipelines. Pairwise sequence alignment is a major contributor to runtimes of many bioinformatics pipelines due to the associated quadratic time complexity. While faster heuristics exist for certain use cases, computing optimal alignments by means of DP is still important for a variety of applications.

We have presented Accelign, a GPU-accelerated open-source pairwise sequence alignment library. At its core it provides a low-level API to high-performance alignment kernel functions for different types of input data, scoring schemes and alignment types. The associated parallelization scheme is based on a common wavefront pattern using warp intrinsics for fast inter-thread communication that minimizes the overall executed instructions using in-register computation in order to unlock the processing power of modern GPU architectures. We provide accelerated kernels for local, global, and semi-global alignment, allowing for the computation of optimal alignment scores with various scoring schemes (linear or affine gap penalties, substitution matrix or PSSM) and alphabets (e.g. DNA/RNA, proteins) supporting one-to-one, one-to-all, and all-to-all use cases on a single or on multiple GPUs. In addition, start/end positions of optimal alignments can be provided. Computing detailed tracebacks is currently not supported, but based on start/end positions highly tuned CPU-based implementations for traceback of global alignments such as BlockAligner could efficiently be used for this task.

Our performance evaluation shows how Accelign can be used to accelerate a number of case studies: short read alignment, protein database scanning, and HIV read mapping. The associated performance evaluations show that it outperforms prior GPU-based libraries (GASAL2, ADEPT) and CPU-based libraries (SeqAn, BSalign, Parasail) as well as specialized GPU implementations (WFA-GPU, CUDASW++4.0).

Our evaluation also demonstrates that speed can further increase by using newer GPU generations and by using multiple GPUs connected to the same host.

While the simple high-level library functions can be used to immediately accelerate applications without additional tuning efforts, the best performance might only be achieved by a custom configuration tuned towards a specific application. Our grid-search was only performed on a subset of possible matrix tile sizes. Another opportunity for improvement lies in the choice of matrix tile size for multi-tile processing depending on the sequence length. Ideally, we would need to consider the actual performance values of different tile sizes, which could be included in future releases. Based on our generic parallelization strategy the library can be easily extended for other DP-based algorithms such as dynamic time warping for Nanopore signals, the Viterbi algorithm for profile HMMs, or profile-to-profile alignments.

## Availability and requirements

Project name: Accelign

Project home page: https://github.com/fkallen/Accelign

Operating system(s): Linux

Programming language: C++, CUDA

Other requirements: CUDA toolkit ≥ 12.9 License: Apache 2.0

Any restrictions to use by non-academics: No additional restrictions

## Declarations

## Ethics approval and consent to participate

Not applicable.

## Consent for publication

Not applicable.

## Availability of data and materials

Accelign is available at https://github.com/fkallen/Accelign. The datasets used for evaluation can be found at https://zenodo.org/records/17865095.

## Authors’ contributions

FK is the main developer of Accelign. FK and FD conducted the experiments. BS proposed and supervised the project. BS and FK wrote the draft of the manuscript. BS prepared figures 1-2. FK prepared figures 3-5. FD prepared figure 6. FK, FD, MS, and BS edited the manuscript. All authors read and approved the final manuscript.

## Acknowledgments

This work was partially funded by Deutsche Forschungsgemeinschaft (DFG, German Research Foundation) – project number 439669440 TRR319 RMaP TP C01 and the NVIDIA Academic Grant Program. M.S. acknowledges support by the Novo Nordisk Foundation (NNF24SA0092560).

## Footnotes

1 https://www.ebi.ac.uk/ena/browser/view/SRR1766553

